# Environmental RNA interference in two-spotted spider mite, *Tetranychus urticae*, reveals dsRNA processing requirements for efficient RNAi response

**DOI:** 10.1101/2020.05.10.087429

**Authors:** Nicolas Bensoussan, Sameer Dixit, Midori Tabara, David Letwin, Maja Milojevic, Michele Antonacci, Pengyu Jin, Yuka Arai, Kristie Bruinsma, Takeshi Suzuki, Toshiyuki Fukuhara, Vladimir Zhurov, Sven Geibel, Ralf Nauen, Miodrag Grbic, Vojislava Grbic

**Affiliations:** Department of Biology, The University of Western Ontario, London, Ontario, N6A 5B8, Canada; Department of Applied Biological Science, Tokyo University of Agriculture and Technology, Fuchu, Tokyo, 183-8509, Japan; Graduate School of Bio-Applications and Systems Engineering, Tokyo University of Agriculture and Technology, Koganei, Tokyo, 184-8588, Japan; Institute of Global Innovation Research, Tokyo University of Agriculture and Technology, Fuchu, Tokyo, 183-8509, Japan; Division Crop Science, Research and Development, Bayer AG, Monheim, Germany; Instituto de Ciencias de la Vid y el Vino, Logrono, 26006, Spain; Department of Biology, University of Belgrade, Belgrade, 11000, Serbia

## Abstract

Comprehensive understanding of pleiotropic roles of RNAi machinery highlighted the conserved chromosomal functions of RNA interference. The consequences of the evolutionary variation in the core RNAi pathway genes are mostly unknown, but may lead to the species-specific functions associated with gene silencing. The two-spotted spider mite, *Tetranychus urticae*, is a major polyphagous chelicerate pest capable of feeding on over 1,100 plant species and developing resistance to pesticides used for its control. A well annotated genome, susceptibility to RNAi and economic importance, make *T. urticae* an excellent candidate for development of an RNAi protocol that enables high-throughput genetic screens and RNAi-based pest control. Here, we show that the length of the exogenous dsRNA critically determines its processivity and ability to induce RNAi *in vivo*. A combination of the long dsRNAs and the use of dye to trace the ingestion of dsRNA enabled the identification of genes involved in membrane transport and 26S proteasome degradation as sensitive RNAi targets. Our data demonstrate that environmental RNAi can be an efficient reverse genetics and pest control tool in *T. urticae*. In addition, the species-specific properties together with the variation in the components of the RNAi machinery make *T. urticae* a potent experimental system to study the evolution of RNAi pathways.

## Introduction

The ability of double-stranded RNA (dsRNA) to inhibit the expression of a complementary target gene in a process named RNA interference (RNAi) was first demonstrated in plants and *Caenorhabditis elegans*^1,2^ and was since discovered in a wide range of organisms^3–5^. The RNAi is guided by small RNAs that can be generated from different precursor RNA molecules. Short interfering RNAs (siRNAs) are generated from dsRNAs. They interfere with the expression of viral- and transposon-originating transcripts or direct chromatin modifications that are essential for proper chromosomal functions^6,7^. Micro-RNAs (miRNAs) are processed from genome-encoded hairpins that contain local stem-loop structures. These precursor RNAs undergo a series of maturation steps to generate Argonaute-associated miRNAs that primarily silence protein-coding mRNAs through either translational inhibition or mRNA degradation. Finally, single-stranded RNA (ssRNA) precursors, transcribed from chromosomal loci that mostly consist of remnants of transposable element sequences, give rise to piRNAs. They are mainly active in germ-line tissues, where they transcriptionally silence transposable elements^7^.

Understanding of the siRNA RNAi pathway led to the use of the exogenously supplied dsRNA to trigger RNAi (termed environmental RNAi^8^). This opened possibilities to develop RNAi as reverse genetics tool^9–13^ and more recently as environmentally friendly strategy for pest control^14,15^. The stability of the exogenous dsRNAs in body fluids, their cellular uptake, and spread to distal tissues that were not in direct contact with the initially delivered dsRNA were identified as key elements for the efficient environmental RNAi. However, the variability in the gut pH and the presence of dsRNases in gut and/or hemolymph that lead to dsRNA hydrolysis and degradation, limit the use of RNAi in many insect clades (reviewed in Cooper *et al.*^16^). Consequently, many arthropods (e.g. some lepidopterans) are refractory to the environmental RNAi. Others (such as some coleopterans, blattodeans and orthopterans) can induce the environmental RNAi only by the injection of dsRNA, restricting its use to reverse genetics. Finally, only a subset of arthropods that respond to the orally delivered dsRNAs are potential candidate species for whom environmental RNAi can be used as a reverse genetics and a pest control tool^16^.

The two-spotted spider mite, *Tetranychus urticae*, is an important agricultural pest that feeds on more than thousand plant species including over 150 crops^17^. Consistent with its outstanding xenobiotic resistance against plant allelochemicals, *T. urticae* populations can rapidly develop resistance to pesticides used for its control^18^, hindering the control of mite infestations in agricultural settings. *T. urticae* is also a model chelicerate whose genome is well annotated and constantly updated^19^. In addition, numerous datasets describing mite transcriptional changes associated with developmental progression, pesticide resistance, or upon host-shift, are available^19–24^. Furthermore, the candidate loci underlying mite physiological processes (e.g. diapause, carotene biosynthesis) and pesticide resistance using forward genetic approaches have been recently identified^25,26^. However, the assessment of the *in vivo* function of candidate loci, at the level of otherwise unperturbed whole organism, is so far lacking. In the absence of the well-established reverse genetics tools, demonstration that proteins encoded by candidate loci have capabilities to modify mite physiology rely on the expression of candidate genes in heterologous systems or on *in vitro* assays. Thus, development of the reverse genetics tools is essential for the understanding of gene function and the unique features of *T. urticae* biology.

The analysis of *T. urticae* genome identified significant variation in RNAi core machinery compared with other arthropods^19^. *T. urticae* harbours a single copy of *pasha* and *drosha* genes, two *dicer* homologues, seven *argonaute* and seven *piwi* genes. *T. urticae* lacks *R2D2* that is a co-factor of Dicer-2 in *Drosophila*, however, it contains two copies of *loquacious* gene. Finally, unlike in insects and crustaceans, mite genome contains five homologs of *RNA dependent RNA polymerase (RdRP)* genes^19^. Pleiotropic roles of RNAi in gene silencing and chromosomal functions (reviewed in Gutbrod and Martienssen^6^), together with the contribution of core RNAi proteins to more than one RNAi pathway, confines an understanding of the functional implications of the diversity and evolutionary variation in RNAi machinery^7,16,27,28^.

Regardless, it has been demonstrated that maternal injections of dsRNA and siRNA, and oral delivery of dsRNA can induce environmental RNAi in *T. urticae*^29–32^. Here, we investigated the effect of sequence composition and length of the exogenous dsRNA on the efficiency of RNAi and its processivity by Dicer using both *in vivo* and *in vitro* analysis. Understanding the length requirement of the exogenous dsRNA and utilization of tracer dye to control the variability in dsRNA ingestion led to development of a highly efficient RNAi protocol. The protocol was rigorously tested against twelve *T. urticae* homologs of *Tribolium castaneum* sensitive RNAi targets^33^, demonstrating that environmental RNAi can be an efficient reverse genetics and pest control tool in *T. urticae*. In addition, the species-specific properties together with the variation in the complement of RNAi pathway genes identified in *T. urticae* genome make the two-spotted spider mite a potent experimental system to study the evolution of RNAi pathways.

## Results

### *TuCOPB2* is a sensitive target for RNAi

We previously identified *TuVATPase* as a sensitive RNAi target. However, only about 50% of treated mites responded to the application of dsRNA and developed RNAi-associated phenotype^32^. To ensure that the improvement of the RNAi protocol is independent of the target gene and results in the general enhancement of RNAi efficiency, we first identified additional target whose silencing leads to trackable phenotype. Previously, Kwon *et al*.^30,31^ identified *TuCOPB2* as a sensitive RNAi target following leaf-disc mediated delivery of dsRNA. To test the ability of dsRNA-*TuCOPB2* to induce RNAi-associated phenotypic changes, we designed two dsRNA-*TuCOPB2* fragments (fragment A of 308 bp and B of 513 bp (Figure 1A)) and delivered them to mites using the soaking method^32,34^. As a control, we used the dsRNA-NC (382 bp in length) that has complementarity against the non-transcribed intergenic region of the *T. urticae* genomic scaffold 12, Figure 1A. Two days post dsRNA treatment, 62% and 51% of mites treated with dsRNA-*TuCOPB2-*A and -B, respectively, exhibited smaller size and a light red-orange color, referred to as a spotless phenotype (Figure 1B; *spotless*). Mites treated with dsRNA-NC invariably displayed the two distinctive black spots (Figure 1B; *normal*). To ensure that the spotless phenotype is associated with the RNAi response, we grouped mites post dsRNA-*TuCOPB2* treatment based on their body phenotype and characterized mite fitness parameters and the expression of the *TuCOPB2* in these mite subpopulations separately. The survivorship and the fecundity of mites treated with dsRNA-*TuCOPB2*-A or -B, displaying the normal body phenotype, were similar to these fitness parameters observed in mites treated with dsRNA-NC (Figure 1C and D; *normal body phenotype*). However, mites treated with dsRNA-*TuCOPB2*-A or -B, displaying the spotless phenotype, had significantly lower survivorship and dramatically reduced fecundity (≥90% reduction) compared to the control (Figure 1C and D; *spotless body phenotype*). Furthermore, the expression level of the endogenous *TuCOPB2* transcripts was significantly reduced (by about 50%) in spotless mites relative to mites treated with dsRNA-NC (Figure 1E). Thus, reminiscent of the effectiveness of dsRNA-*TuVATPase*, dsRNA-*TuCOPB2* induced strong RNAi in approximately 50% of treated mites. The silencing of *TuCOPB2* resulted in a trackable phenotype, identifying an independent target gene that can be used together with *TuVATPase* for the improvement of RNAi efficiency in *T. urticae*.

**Figure 1.**
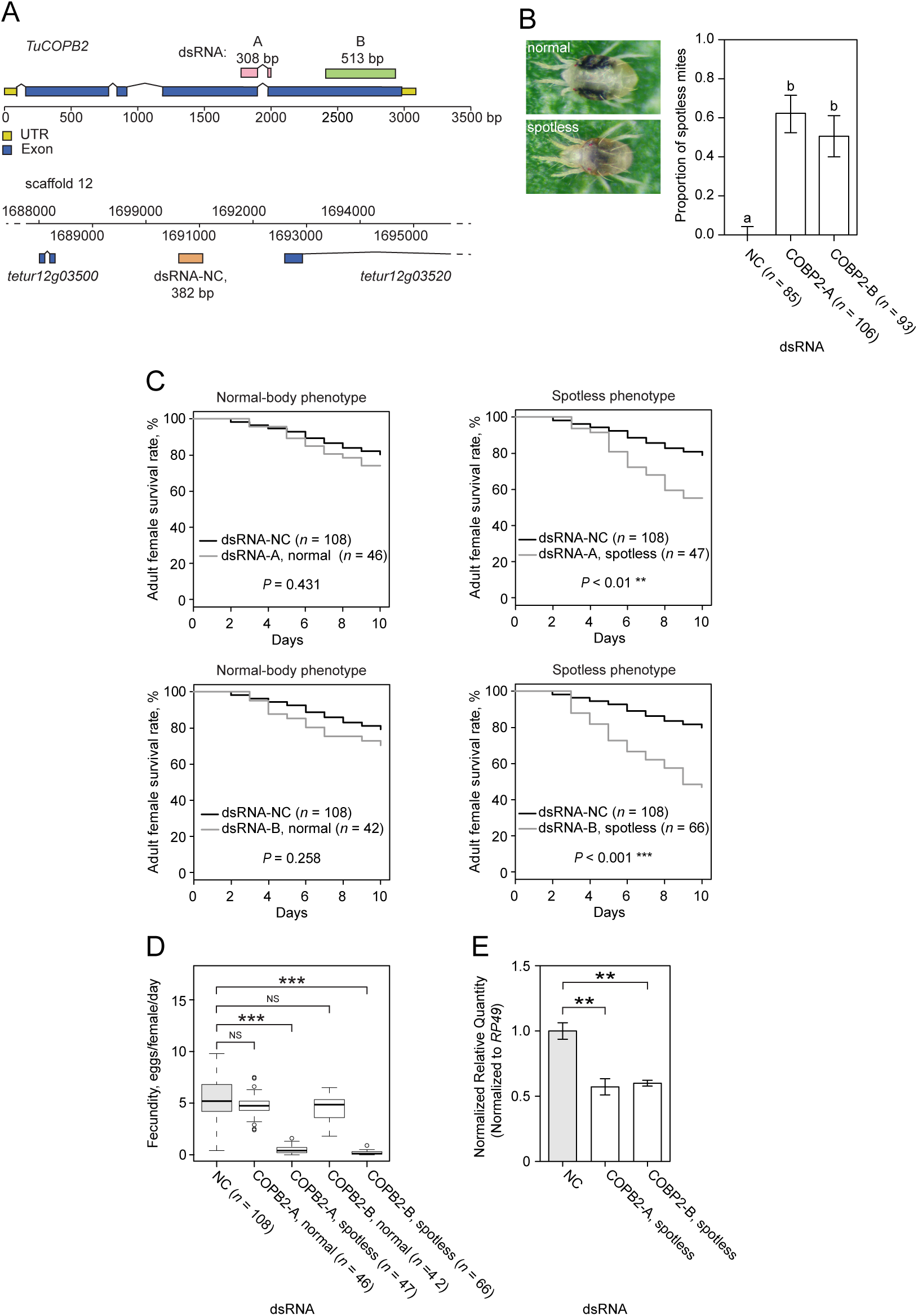
dsRNA-*TuCOPB2* induces a spotless body phenotype in a portion of treated adult female mites. (**A**) Schematics of the *TuCOPB2* locus depicting DNA sequences used for the generation of dsRNA-*TuCOPB2-*A and dsRNA-*TuCOPB2-*B. The part of the scaffold 12 of *T. urticae* genome depicting the location of the 382 bp fragment that was used to synthesize dsRNA-NC. (**B**) Female mites showing the normal and spotless phenotypes, and the proportion of mites displaying the spotless phenotype after soaking mites in the solution of dsRNA-NC, dsRNA-*TuCOPB2-*A, and dsRNA-*TuCOPB2*-B (160 ng/μL). The bars represent the proportion of mites displaying phenotype and error bars represent 95% CI of the proportion. Data were analyzed with Fisher’s exact test followed by Bonferroni correction for multiple comparisons. Different letters indicate significant differences, *P*<0.05. (**C**) Adult mite survivorship over 10 days after soaking in solution of *TuCOPB2-*A, dsRNA-*TuCOPB2*-B, or dsRNA-NC (160 ng/μL). Survival curves of normal and spotless mites were plotted using the Kaplan-Meier method and compared using the log-rank test with Bonferroni correction (**, *P*<0.01; ***, *P*<0.001). (**D**) Fecundity of normal and spotless mites after soaking in the solution of *TuCOPB2-*A, dsRNA-*TuCOPB2*-B or dsRNA-NC. Statistical analysis was performed using the Wilcoxon-Mann-Whitney test with Bonferroni correction (*, *P*<0.05; ***, *P*<0.001). (**E**) *TuCOPB2* expression levels were normalized to the expression of the reference gene *RP49* and were shown relative to the expression in the negative control (NC). Data are represented as a mean ± SEM. Data were analysed with one way-ANOVA followed by Dunnett’s test (**, *P* < 0.01). All experiments were performed in 3 independent experimental trials.

### Long dsRNAs are required for the efficient RNAi in *T. urticae*

The length of dsRNAs has been shown to be an important parameter affecting the efficiency of RNAi in a wide range of organisms^35–38^. To test the effect of dsRNA size on RNAi efficiency in *T. urticae*, a series of nested dsRNA fragments against *TuVATPase* and *TuCOPB2* were evaluated for their ability to induce RNAi-associated mite phenotypes (dark-body phenotype upon the application of dsRNAs targeting the *TuVATPase* and spotless phenotype upon the application of dsRNAs against the *TuCOPB2*), Figure 2A. The longest fragment had 600 bp (shown in red), while shorter fragments (400, 200 and 100 bp) were overlapping at either the 5’ (shown in green) or the 3’ (shown in blue) ends of the 600 bp-fragment. The effectiveness of dsRNAs fell in two classes: a) short dsRNAs, 100 and 200 bp in length, induced limited RNAi responses (≤4.4% and ≤19% of treated mites developed an RNAi-associated phenotypes when treated with 100 bp and 200 bp dsRNA fragments, respectively); and b) dsRNA fragments of 400 and 600 bp had similar RNAi efficiency and induced RNAi-associated phenotypes in ≥46% of treated mites, Figure 2A.

**Figure 2.**
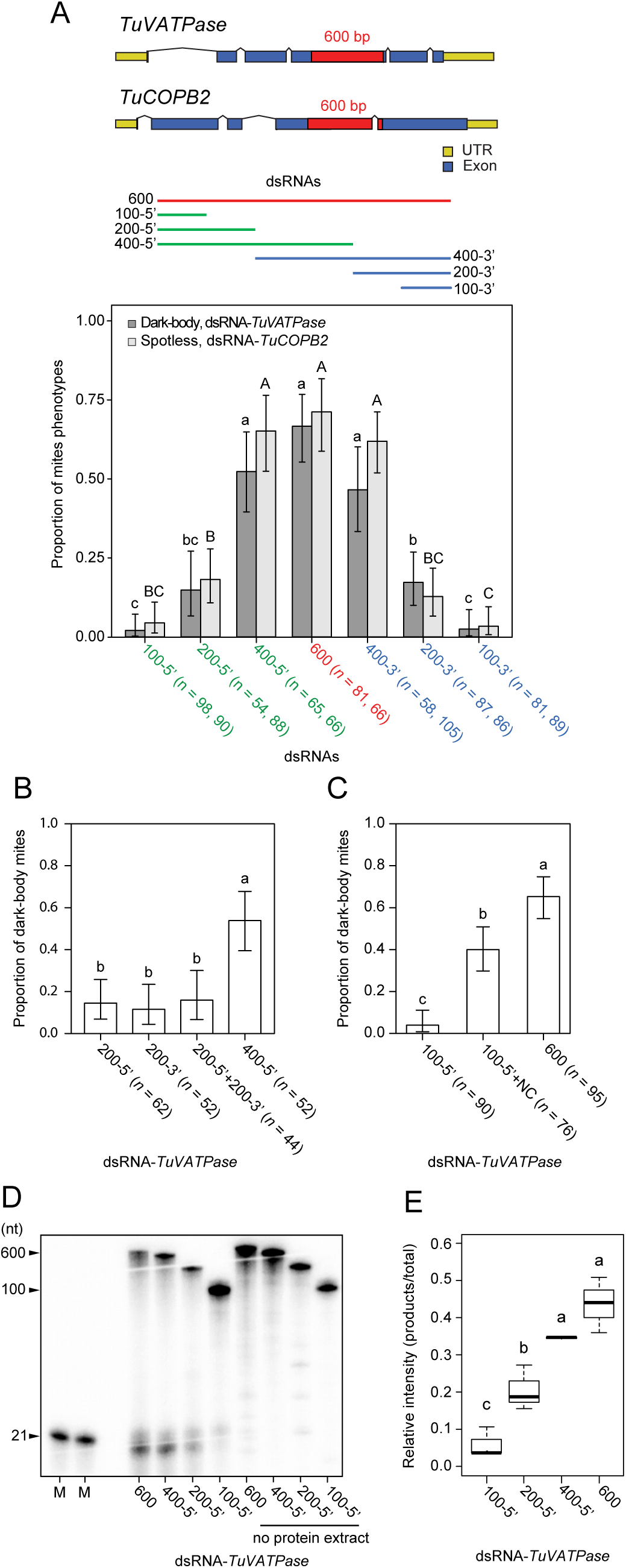
Long dsRNAs are required for the efficient processing of dsRNAs and RNAi. (**A**) The effect of dsRNAs length on RNAi efficacy. Sizes and positions of dsRNAs targeting *TuCOPB2* and *TuVATPase* are shown in red for the longest dsRNAs (600 bp), in green for the shorter overlapping fragments (400, 200 and 100 bp) positioned at the 5’ ends of the 600 bp-fragment, and in blue for the shorter fragments positioned at the 3’ ends of the 600 bp-fragment. RNAi efficacy is measured as the frequency of spotless (after treatment with dsRNA-*COPB2*) and dark-body (after treatment with dsRNA-*TuVATPase*) phenotypes. (**B**)Frequency of dark mites upon the application of dsRNAs of different lengths and sequence coverage. dsRNA-*TuVATPase* fragments spanning 200 bp at either the 5’ or 3’ end of the 600 bp-fragment were applied individually or as a mix. A 400-5’ bp dsRNA-*TuVATPase* fragment was used as a control. (**C**) Frequency of dark mites upon treatment with the dsRNA-*TuVATPase* 100-5’ and the chimeric dsRNA fragment that is composed of 100-5’ bp *TuVATPase* fragment and 385 bp fragment corresponding to dsRNA-NC sequence. Application of dsRNA-*TuVATPase* 600 bp was used as a control. (A-C) All fragments were applied at 160ng/μL. Data represent the 95% confidence interval of the proportion of mites displaying phenotype. Data were analyzed with Fisher’s exact test followed by Bonferroni correction for multiple comparisons. Different letters indicate significant differences, *P*<0.05. (**D**) Dicing (dsRNA-cleaving) activities of crude mite extracts incubated with ^32^P-labeled dsRNA-*TuVATPase* of different lengths (600, 400, 200, and 100 bp) at 22°C for 2 h. Long dsRNAs and processed small RNAs were detected by autoradiography. No crude extract was used as the negative control. M indicates a size marker (21 nt). (**E**) The radioactive intensity of siRNA products relative to the total intensity. Data were analyzed with one-way ANOVA followed by Tukey’s HSD test. Different letters indicate significant differences, *P*<0.05. All data were collected from three independent experimental runs.

The greater efficiency of longer dsRNA fragments could be due to greater target sequence coverage of siRNAs generated from dsRNA precursor or can arise from a greater efficiency of cellular uptake and/or processing. To differentiate between these possibilities, we first compared the RNAi efficacy of a mix of two dsRNAs of 200 bp in length relative to the application of a single dsRNA of 400 bp. If uptake and processing of these dsRNAs are similar, then these treatments are expected to have similar RNAi efficiency as they are expected to generate similar complement of siRNAs. However, if the uptake and/or processing of dsRNAs of different lengths differ, the RNAi efficacy of long dsRNA is expected to be greater than that of short dsRNAs. As shown in Figure 2B, dsRNAs of 200 bp in length, either individually or as a mix, had significantly lower ability to induce RNAi-associated phenotype relative to the 400 bp fragment, suggesting that ineffectiveness of short dsRNAs is not due to the lack of sequence variety. To further understand the contribution of dsRNA length on the effectiveness of RNAi, we prepared the chimeric dsRNA fragment consisting of a 100-5’ sequence complementary to *TuVATPase* (Figure 2A) and 382 bp of non-transcribed sequence used in the dsRNA-NC (Figure 1A). Comparison of RNAi responses triggered by the 100-5’dsRNA-*TuVATPase* and the chimeric fragment revealed that the chimeric fragment was about 11 times more efficient, Figure 2C. As these fragments are expected to give rise to similar pools of effective siRNAs, the differential effectiveness of these dsRNAs indicates that their uptake or processivity may differ.

To directly test if the processivity of dsRNAs is dependent on the length of dsRNAs, we established a biochemical method to monitor the Dicer dsRNA-cleaving activity in mite cell-free extracts. By mixing the crude mite protein extract with ^32^P-labeled dsRNAs of different lengths as substrates, we were able to visualize the cleaved RNA products using denaturing polyacrylamide gel electrophoresis (PAGE) and autoradiography. As seen in Figures 2D and E, ^32^P-labeled small RNAs tightly correlated with the progressive increase of dsRNA length (Figure 2E). These results demonstrate that Dicer preferentially cleaves long dsRNAs in *T. urticae*. Thus, dsRNAs of ≥400 bp in length are required for the efficient processing of dsRNAs and RNAi in *T. urticae.*

### Tracing the dsRNA ingestion increases RNAi responsiveness to >90% in treated mite populations

Even the length-optimized dsRNAs induced RNAi only in a subset of treated mites, Figure 2. A variation in dsRNA uptake, tracked by the addition of dye into dsRNA solution, was previously reported for aphids^39^. To test if mites soaked in dsRNA solution ingest dsRNA differentially, we used a 6% blue food dye that can be easily visualized in mite posterior midgut, Figure 3A, and mixed it in dsRNA solution. Upon soaking, 64% and 66% of the total mites treated with dsRNA-*TuVATPase*-B and dsRNA-*TuCOPB2*-B showed blue color in posterior midgut, respectively. When mites were preselected for the presence of blue dye in their gut, the proportion of mites displaying the RNAi-associated phenotypes was greater than 90% (94% and 95% of mites treated with dsRNA-*TuVATPase*-B and dsRNA-*TuCOPB2*-B displayed a dark-body and a spotless phenotypes, respectively (Figure 3B)). Mites without the visible accumulation of dye had a significantly lower frequency of body phenotypes associated with RNAi responses (Figure 3B). Thus, the blue dye could be used as a dsRNA tracer, securing the RNAi responsiveness in the majority of treated mites.

**Figure 3.**
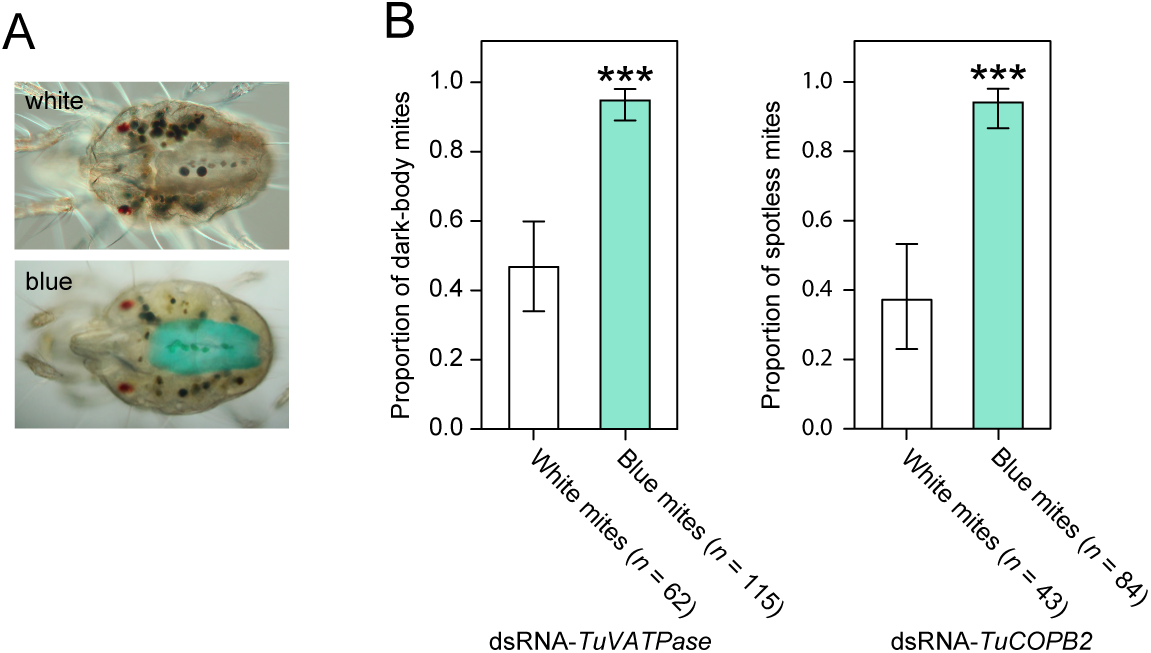
Blue dye can be used as a tracer to correct for the variability in dsRNA ingestion. (**A**) Representative images showing absence (white) or accumulation of tracer dye (blue) in the posterior midgut after mite treatment with a solution containing 6% blue dye. (**B**) Frequency of dark and spotless mites upon mite soaking in the solution of blue dye that was mixed with dsRNA-*TuVATPase* or dsRNA-*TuCOPB2*, respectively. Data represent the 95% confidence interval of the proportion of mites displaying dark-body (on the left) and spotless (on right) phenotypes. Statistical analysis was performed using Fisher’s exact test. ***, *P*<0.001.

### The effect of dsRNA concentration on RNAi efficacy

Several application methods for the oral delivery of dsRNA to *T. urticae* have been developed^30,32,38^. These methods use a wide range of dsRNA concentrations, from 40 ng/μL up to 1 μg/μL. To determine the effective dsRNA concentrations and the potential toxicity of high concentrations of dsRNAs due to the possible oversaturation of the RNAi machinery, we applied dsRNA-*TuVATPase*, dsRNA-*TuCOPB2* and dsRNA-NC in the range of 20 to 1280 ng/μL and scored mite survivorship, Figure 4. At 20 ng/μL, individual dsRNAs against *TuVATPase* and *TuCOPB2* did not affect mite mortality, Figure 4A. However, the mixture of these dsRNAs (at the concentration of 20 ng/μL each) acted synergistically and resulted in a significant decrease of mite survivorship, Figure 4B. dsRNA-*TuVATPase* and dsRNA-*TuCOPB2* induced significant mite mortality (80%) at 40 and 80 ng/μL, respectively, and their responses were saturated at 160 ng/μL, Figure 4A. Therefore, even though the effective concentrations of dsRNA may vary dependent on the target, the concentration of 160 ng/μL of dsRNA appears to be sufficient to induce the full RNAi response and was used in subsequent experiments. High concentrations of dsRNAs, at 1280 ng/μL, did not affect mite mortality (Figure 4C), indicating that dsRNA at this concentration was not toxic to mites and did not oversaturate the RNAi machinery.

**Figure 4.**
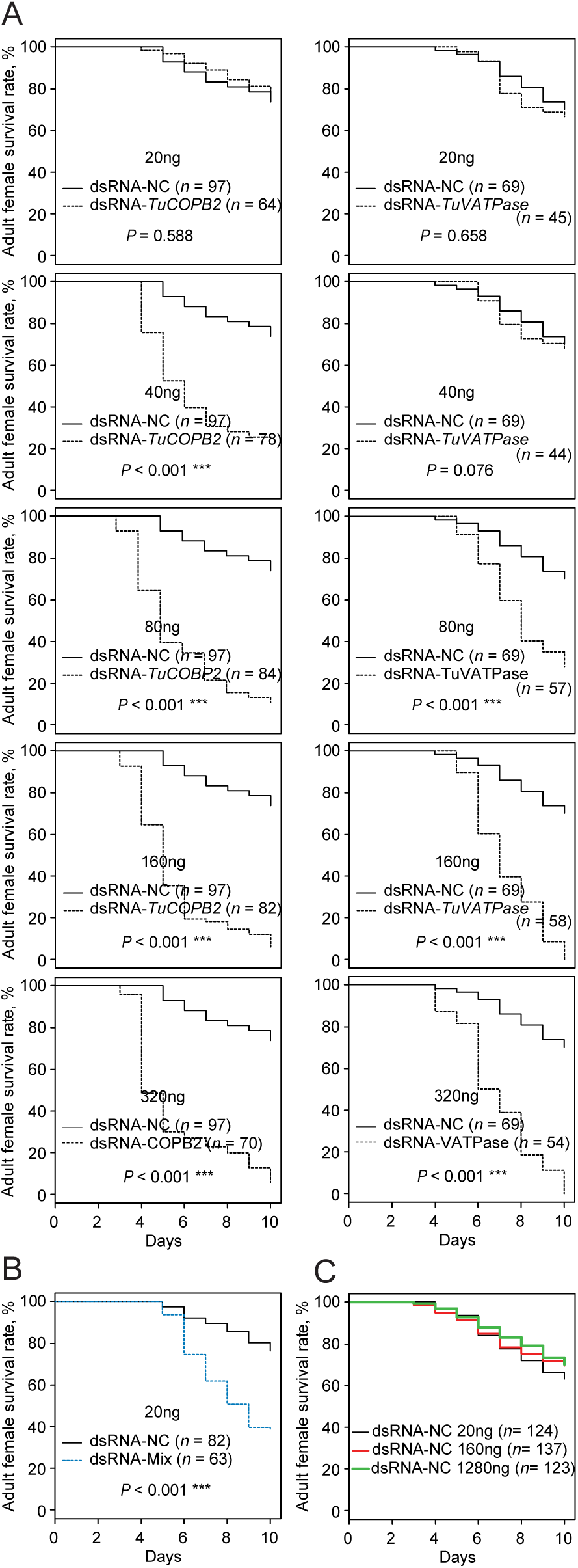
RNAi dose-response curves following dsRNA treatment. (**A**) Adult survivorship following mite treatment with dsRNA-*TuVATPase* and dsRNA-*TuCOPB2* at 20, 40, 80, 160, and 320 ng/μL. (**B**) Synergistic effect of combined application of dsRNA-*TuVATPase* and dsRNA-*TuCOPB2* at 20 ng/μL each, on adult survivorship. (**C**) Adult survivorship following treatment with dsRNA-NC at 20, 160, and 1280 ng/μL. All experiments were performed in 3 independent trials. Survival curves were plotted using the Kaplan-Meier method and compared using the log-rank test with Bonferroni correction (not significant, *P*>0.05; ***, *P*<0.001).

### Testing the effectiveness of RNAi on a broad range of gene targets

Almost complete RNAi responsiveness of mite populations to dsRNA-*TuVATPase* and dsRNA-*TuCOPB2* using the optimized method raises a possibility to use RNAi as a reverse genetics tool in *T. urticae*. However, the use of RNAi as a tool to induce loss-of-function phenotypes requires a demonstration of RNAi effectiveness on a broader range of target genes. Previously, Ulrich *et al*.^33^ performed an RNAi screen of 5,000 randomly selected *Tribolium castaneum* genes and identified eleven highly efficient RNAi targets whose silencing led to 100 % mortality of *Tribolium* larvae. These genes are predicted to encode conserved proteins that perform essential cellular functions involved in membrane transport, degradation of ubiquitinated proteins through 26S proteasome, protein folding and secretion, and regulation of gene expression, Table 1, and were proposed to be conserved candidate RNAi targets across arthropods. Thus, to test the efficiency of the optimized RNAi protocol on a greater number of genes, we identified *T. urticae* orthologs of *Tribolium* RNAi targets, Table 1. The search of *T. urticae* orthologs revealed that *T. urticae* genome contains two copies of *Pp1a-96a* locus, both of which were included in our analysis. *Pp1a-96a* orthologs share 78% CDS sequence identity (95% aa identity) and are further referred to as *TuPp1α-96a-A* and *TuPp1α-96a-B*.

**Table 1:**
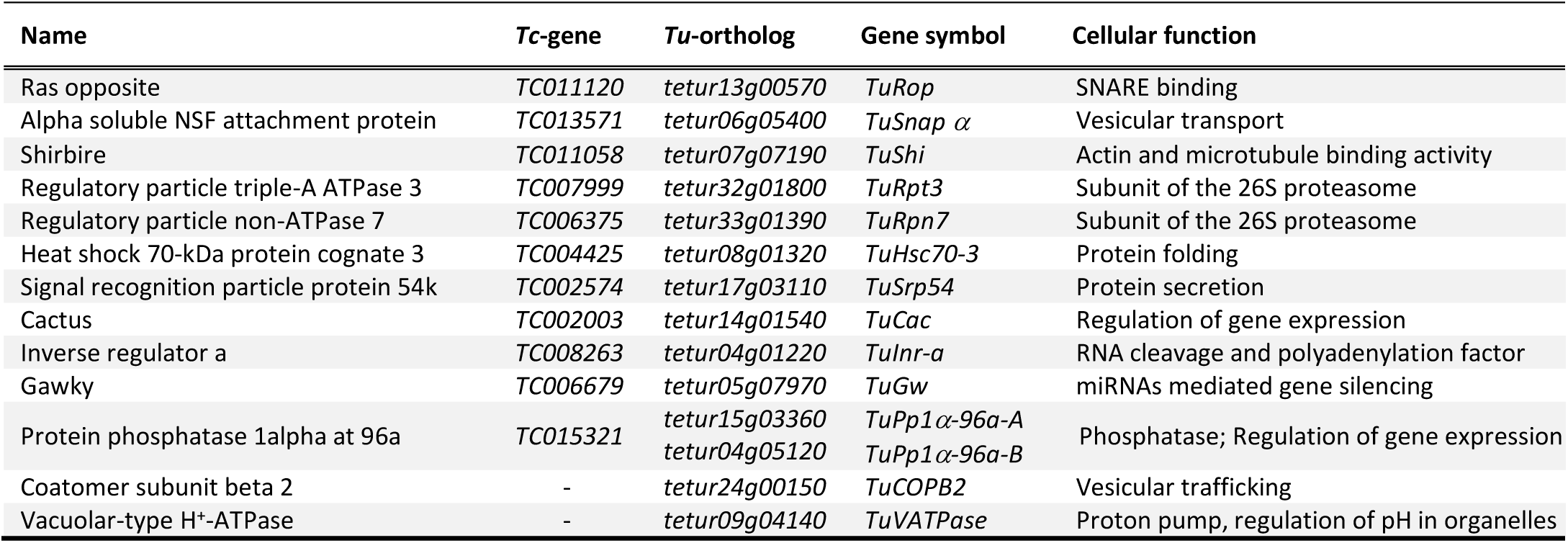
List of *Tetranychus urticae* orthologues of *Tribolium castaneum* genes identified as highly efficient RNAi targets

dsRNA treatments targeting *TuRpn7, TuSnap α, TuRop*, and *TuSrp54* resulted in a dark-body phenotype previously observed in dsRNA-*TuVATPase* treated mites, Figure 5. In addition, mites treated with dsRNAs against *TuRpt3* and *TuHsc70-3* displayed a spotless phenotype characteristic for mites treated with dsRNA-*TuCOPB2*. dsRNAs against *TuInr-a, TuGw, TuShi, TuPp1α-96a-A*, and *TuPp1α-96a-B* yielded mite populations whose phenotypes were indistinguishable from mites treated with dsRNA-NC, Figure 5 and not shown. The body phenotypes were visible 2 days post-treatment, except for mites treated with dsRNAs against *TuRpt3* and *TuHsc70-3* that required 4 to 5 days post dsRNA treatment for the establishment of the body-color change. The appearance and the timing of body color phenotypes were the same upon application of two independent dsRNA fragments, Supplemental Table 1, indicating that these phenotypes are associated with the RNAi of their corresponding targets.

**Figure 5.**
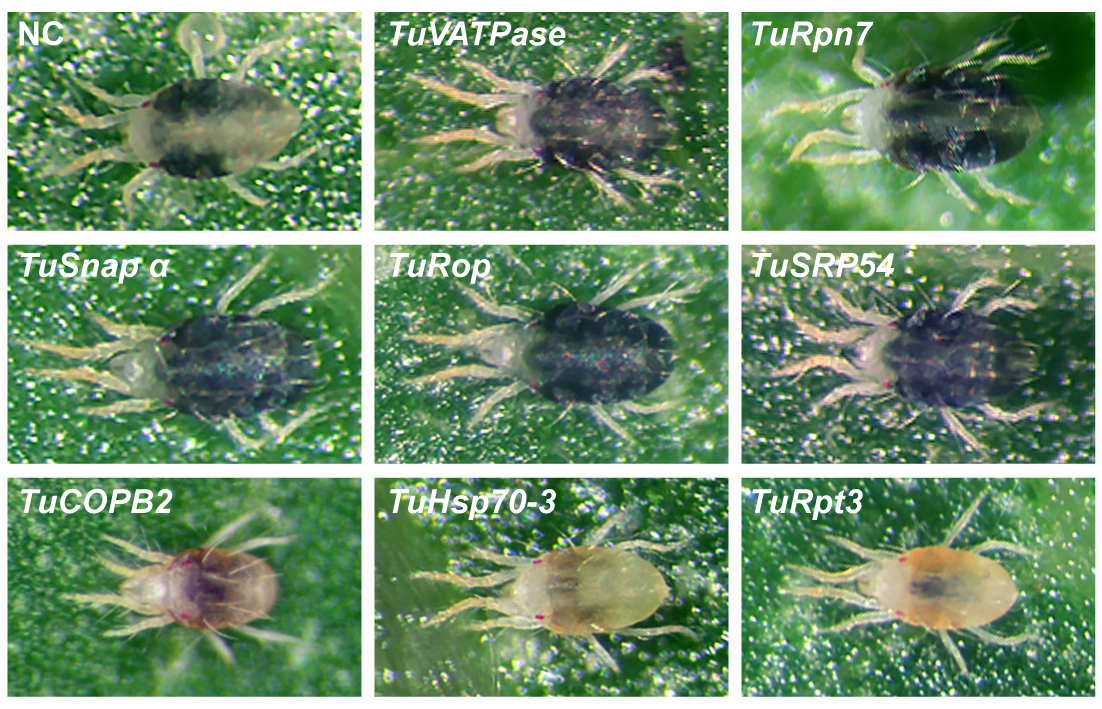
Mite RNAi-associated phenotypes upon the application of dsRNA against *T. urticae* orthologs of *Tribolium* RNAi sensitive targets. Gene targets are listed at the upper left corner. NC, mites were soaked in the solution of dsRNA-NC.

To assess the efficiency of RNAi upon application of these dsRNAs, we determined mite survivorship over 10 days (Figure 6 and Supplemental Figure 1) and fecundity over 3 days (Figure 7) post dsRNA treatments. Even though silencing of all tested target genes resulted in 100% mortality in *Tribolium*^33^, in *T. urticae*, their effectiveness was variable. dsRNAs targeting *TuSrp54, TuRop, TuSnap α, TuRpn7, TuRpt3, TuInr-a* and *TuGw* caused *>*80% mortality of adult female mites, Figure 6. However, mites treated with dsRNA-*TuShi*, dsRNA-*TuHsc70-3*, and a mixture of dsRNAs against *TuPp1α-96a-A* and *TuPp1α-96a-B* had a modest increase in mortality (∼20% more than mites treated with the dsRNA-NC that was used as a negative control), while dsRNA targeting *TuCac* and *TuPp1α-96a-A* and *TuPp1α-96a-B* genes individually had no effect on mite survivorship, Figure 6. Reproducibility of RNAi phenotypes upon the application of a second non-overlapping dsRNA for each gene rigorously confirms that the observed phenotypes are specific to the RNAi of target genes, making off-target effects improbable (Supplemental Figure 1).

**Figure 6.**
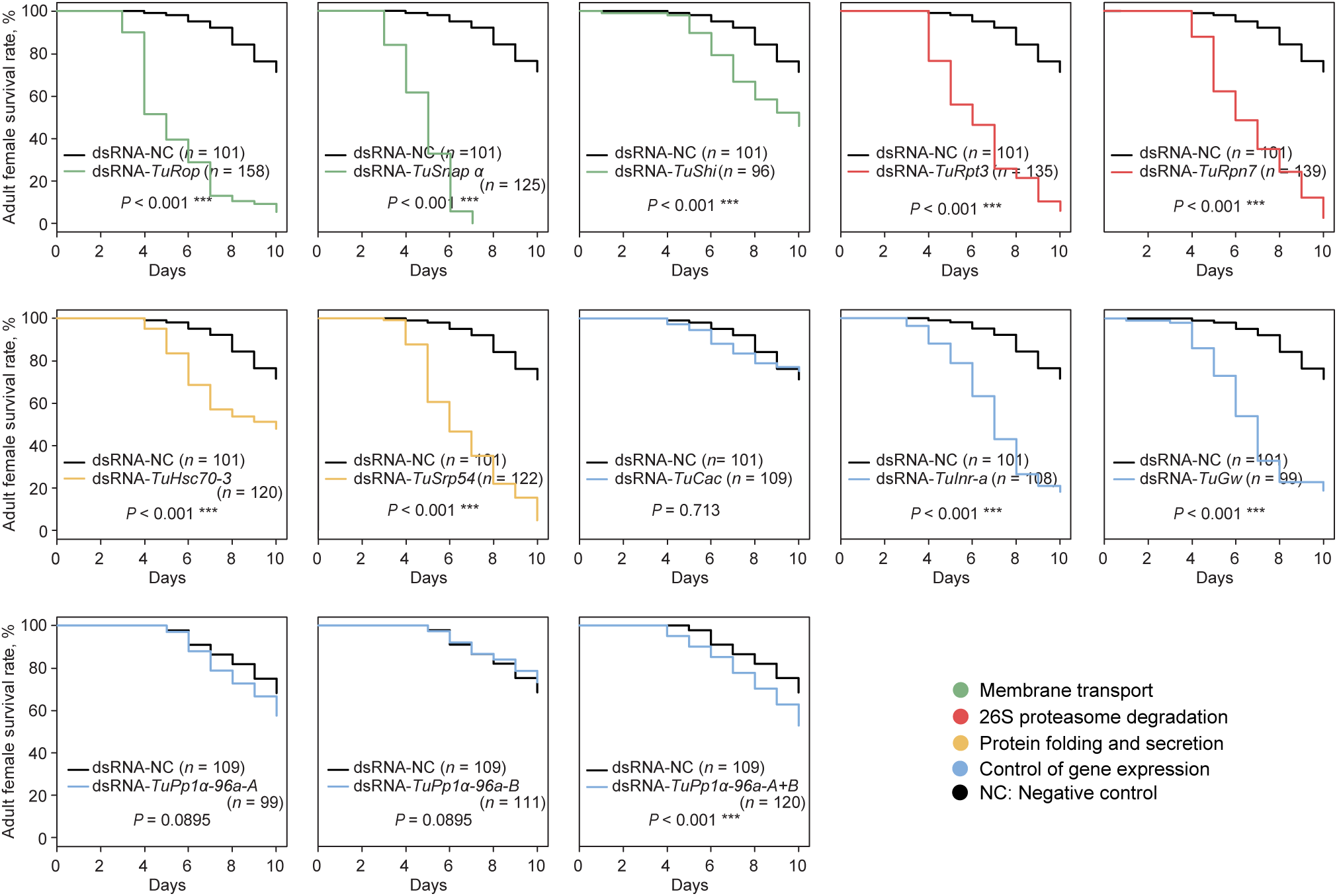
Survival curves of adult female mites after treatment with dsRNAs targeting *T. urticae* orthologs of *Tribolium* RNAi sensitive genes. dsRNAs were applied at 160ng/μL. Survival curves were plotted using the Kaplan-Meier method and compared using the log-rank test with Bonferroni correction (not significant, *P*>0.05; ***, *P*<0.001). All experiments were performed in 3 independent trials.

**Figure 7.**
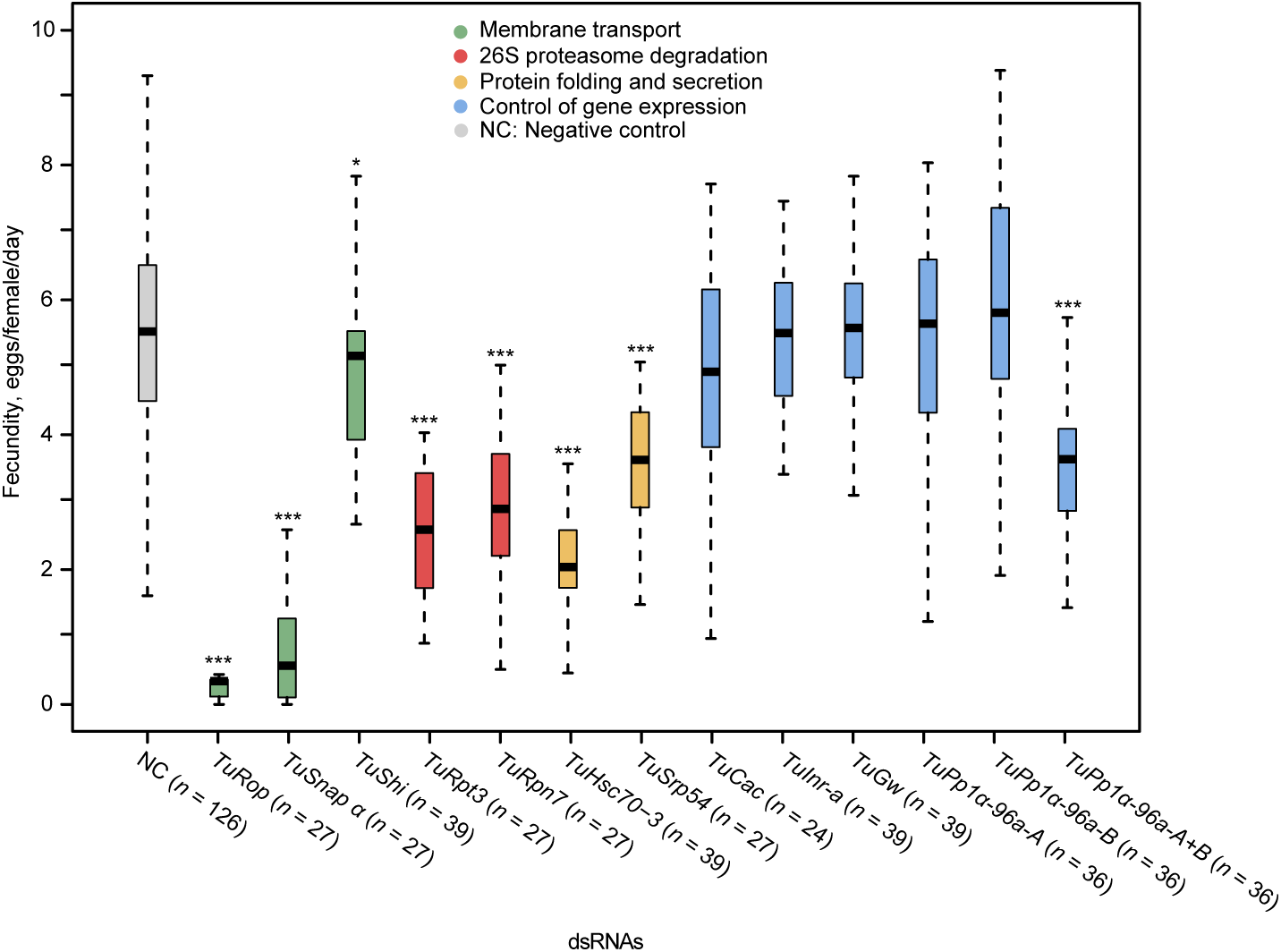
Fecundity of adult female mites after treatment with dsRNAs targeting *T. urticae* orthologs of *Tribolium* RNAi sensitive genes. dsRNAs were applied at 160ng/μL. Fecundity is represented as the average number of eggs per female per day laid over 3 days. Significant differences relative to dsRNA-NC were determined by repeated-measures ANOVA followed by pairwise comparisons of treatment estimated marginal means with a Tukey p-value adjus™ent. (not significant, *P*>0.05; *, *P*<0.05; ***, *P*<0.001). All experiments were performed in 3 independent trials.

Besides mortality, mite fecundity was also affected by the application of dsRNA, Figure 7. Mites treated with dsRNA against *TuRop* and *TuSnap α* had dramatically reduced fecundity, laying on average 0.4 and 0.7 eggs/female/day, relative to mites treated with dsRNA-NC that laid 5.2 eggs/female/day. Most other treatments resulted in ∼50% reduction of fecundity, while dsRNAs targeting *TuShi, TuInr-a, TuGw, TuPp1α-96a-A*, and *TuPp1α-96a-B* did not significantly affect mite egg deposition. Overall, our data demonstrate that RNAi is effective to silence a range of gene targets in *T. urticae*.

## Discussion

The use of RNAi as a reverse genetics tool requires efficient silencing of the expression of the target gene(s) and the responsiveness of the majority of treated individuals. In addition, the ability of ingested dsRNA to silence the target pest gene, resulting in pest mortality or reduced fitness, is a prerequisite for the development of RNAi-based pest control strategies. Many economically important lepidopteran, orthopteran, blattodean, and some coleopteran pests are recalcitrant to RNAi silencing by feeding that impedes the development of RNAi-based tools for their control^16^. The ability of orally delivered dsRNA to induce RNAi in *T. urticae* has been previously established^30–32^, making this species a potential model for the development of RNAi-based pest controls. In this work, we optimized the dsRNA design and tested the utilization of dye as a marker for dsRNA ingestion. These modifications resulted in almost 100% RNAi efficiency. Using the optimized protocol, we demonstrated that sensitive RNAi targets identified in one organism are not directly transferable to another. Rather, the identification of RNAi targets has to be species-specific. The optimized RNAi protocol developed and used in this study, in conjunction with high quality *T. urticae* genome, paves the way for the high-throughput genetic analysis of gene function, for the identification of RNAi targets for mite pest control and for dissection of RNAi processing machinery in this model pest.

The power of RNAi as a reverse genetics strategy is best illustrated by the genome-wide RNAi screens performed in *Tribolium castaneum*^12^, *Drosophila melanogaster*^11^, and *Caenorhabditis elegans*^9,10^. The optimized RNAi protocol used in this study can be scaled up and used for the whole-genome functional analysis of *T. urticae* genes. Such screen would be particularly useful in identifying RNAi target genes for mite pest control, deciphering the function of >4k orphan genes that are unique to *T. urticae*/Chelicerates or the analysis of candidate genes that were identified in studies of mite-plant host interactions^19–24^ and pesticide resistance^26,40^. Design of the gene-specific dsRNA is facilitated by the high quality of *T. urticae* genome sequence and high RNAi efficacy of chimeric dsRNAs, where only a small portion of dsRNA sequence produces effective siRNAs (Figure 2C). However, to claim the loss-of-function phenotype to a target gene and to eliminate the possibility of the off-target RNAi effects, a demonstration that second non-overlapping dsRNA causes the same phenotype is required, Supplemental Figure 1. Furthermore, the ability of a mixture of dsRNAs to simultaneously induce RNAi in several genes enables knock-down of multiple processes (e.g. silencing of *TuVATPase* and *TuCOPB2*, Figure 4B), and the analysis of genes that act redundantly (e.g. *TuPp1α-96a-A* and *TuPp1α-96a-B*, Figures 6 and 7). The CRISPR-Cas9 system is an alternative reverse genetics method for the generation of the loss-of-function phenotypes. The initial demonstration of CRISPR-Cas9 in *T. urticae* has been recently reported^41^, however, it will require a substantial increase in method efficiency to be useful in the generation of mutant alleles. Even when the CRISPR-Cas9 method becomes available, the RNAi approach may be preferred for the generation of loss-of-function phenotypes. RNAi protocols are easy to perform and do not require maintenance of mutant populations. RNAi enables testing the effects of several genes simultaneously simply by combining multiple dsRNAs, which is particularly beneficial in pathway analysis. In addition, RNAi induced decreased gene function can be monitored in a time and dose-dependent manner. However, the other genome editing capabilities of CRISPR-Cas9, e.g. generation of larger deletions or gain-of-function alleles, will be important additional tools for understanding gene functions in *T. urticae*.

The ability of the ingested dsRNA to trigger RNAi in *T. urticae* opens a possibility to develop RNAi-based strategies for its control. As mites easily develop resistance to chemical pesticides^18^, the RNAi-based controls that have an independent mode of action relative to the existing pesticides could be an important tool for mite pesticide resistance management. Here, we tested the ability of twelve *T. urticae* orthologs of *Tribolium* genes that were identified as highly efficient RNAi targets^33^ for their ability to induce RNAi and reduce mite fitness. Though not all candidate genes were confirmed as sensitive RNAi targets in *T. urticae*, silencing of 9 out of 14 genes resulted in a highly significant reduction in fecundity and survivorship (mortality ≥75%), Figures 6-8. These genes represent processes and protein complexes that are particularly susceptible to environmental RNAi in *T. urticae*. Highly effective RNAi targets in *T. urticae* are involved in membrane transport and 26S proteasome degradation of ubiquitinated proteins, Figure 8. In addition, targets involved in the general control of gene expression, *TuGw* (required for gene silencing mediated by miRNAs) and *TuInr-a/Pcf11* (required for the transcriptional termination and 3′ end processing), as well as *TuSRP54* (required for protein secretion) were also sensitive to RNAi in *T. urticae*, Figure 8. In addition to these processes, Kwon *et al.*^30,31^ identified the aquaporins (AQPs) and the regulation of the water balance as the essential process that is sensitive to manipulation by RNAi in *T. urticae.* The high level of mortality and decreased fecundity in mites treated with dsRNAs targeting *TuSNAP α, TuRop* and *TuCOPB2* transcripts (Figures 1, 4 and 8), predicted to dramatically reduce the size of a *T. urticae* population, demonstrate that these or similar RNAi targets can be developed as a tool for mite pest control in agriculture. In addition, dsRNAs targeting mite pesticide resistance genes were demonstrated to act as synergists to the existing pesticides^42–45^ – prolonging their efficiency against mite populations. Identification of additional sensitive gene targets, an understanding of the gut environment that can affect dsRNA stability, as well as mechanisms of dsRNA cellular uptake and spread to target tissues are required for the development of effective RNAi-based crop protection strategies against *T. urticae*. Although these processes are not limiting the efficacy of environmental RNAi in *T. urticae*, they could be the basis of development of mite resistance to dsRNA. For example, the impairment of luminal uptake of dsRNA led to development of dsRNA-resistant western corn rootworm^46^. In insects, SID-1 and SID-1-like (SIL) proteins that encode transmembrane channel proteins, and receptor-mediated clathrin-dependent endocytosis^16,47–49^ were implicated in the cellular uptake of dsRNA. The dsRNA cell import has not been investigated in *T. urticae*. However, in the absence of hemolymph^50^, dsRNA uptake from the gut lumen and transporters that enable intercellular dsRNA movement are expected to profoundly affect the efficiency and the spread of the RNAi in *T. urticae*.

**Figure 8.**
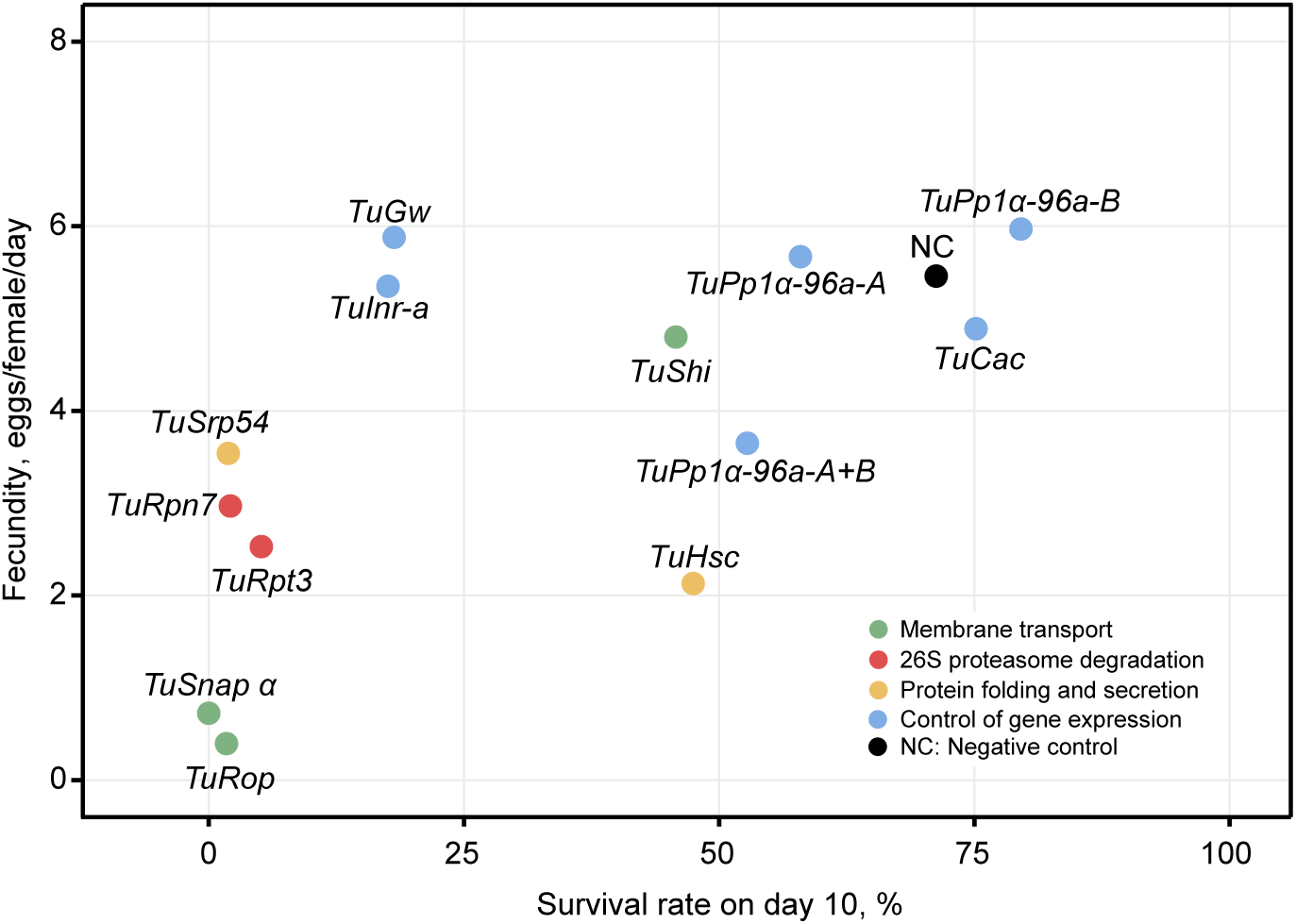
Relationship between mite survivorship rate and fecundity for different classes of target genes. Mite survival rate on day 10 is as shown in Figure 6. Mite fecundity is the mean number of eggs per female per day laid over 3 days as shown in Figure 7.

An understanding of the evolutionary variation in the core RNAi machinery is an important factor in the development of RNAi-based tools. Gene losses and gains, and mutations in individual genes encoding proteins of the core RNAi machinery are expected to lead to species-specific modifications of RNAi pathways. For example, core proteins of the siRNA RNAi pathway include Argonaute-2, Dicer-2, and R2D2. Many beetles have duplicated *R2D2* gene that is associated with efficient environmental RNAi^27^. However, *T. urticae* lacks the homologue of *R2D2* but is still able to induce efficient RNAi by the exogenously supplied dsRNA. Furthermore, the core proteins of the miRNA pathway in insects include Argonaute-1, Dicer-1, Loquacious, Drosha, and Pasha^51^. In *T. urticae*, this pathway is expanded and contains two *dicer* homologues (with unclear homology to *Drosophila dicer*-1 or -2 genes), seven *argonaute* genes, *pasha, drosha* and two copies of *loquacious* gene^19^. Finally, unlike in insects and crustaceans, the *T. urticae* genome contains five homologs of *RNA dependent RNA polymerase (RdRP)* genes that in *C. elegans* are implicated in systemic RNAi response^52^. As the conserved function of RNAi machinery is in the modulation of chromosome segregation, dosage response, and repair of DNA damage^6^, these changes in gene complement may reflect the variations in the generation and gene silencing function of small RNAs. Indeed, we showed that short dsRNAs (≤200 bp) had low ability to induce the RNAi-associated phenotypes and that long dsRNAs (≥400 bp) are required for the efficient RNAi in *T. urticae*. A similar relationship between the length of the dsRNA and RNAi efficacy was observed in other organisms^35,36,53,54^, however, the length of dsRNA required for efficient RNAi differs. For example, unlike in *T. urticae*, dsRNAs of 100-200 bp in length induce highly efficient RNAi in *Tribolium*^36,55^, western corn rootworm^53,54^, *Drosophila* S2 cells^47^ and the nematode *C. elegans*^35^. However, while in *T. urticae* the *in vivo* effects of dsRNAs of different length on RNAi efficacy tightly correlated with the processivity of dsRNA by Dicer *in vitro* assay, Figure 2D and E, in *Drosophila* S2 cells, the length of dsRNA affected its cellular uptake and not the processivity^47^, indicating differences in the RNAi machinery. With the establishment of an efficient RNAi protocol and the identification of sensitive RNAi targets, we are now in position to dissect and characterize RNAi machinery in *T. urticae* and identify evolutionary changes in RNAi genetic pathways that are critical for efficient environmental RNAi.

In summary, we characterized the properties of dsRNA that are essential for the efficient environmental RNAi and demonstrated that RNAi can be used as a reverse genetics and pest control tool in *T. urticae*. In addition, we identified a causality between the length of dsRNA and its processivity, that warrants further dissection of RNAi processing machinery in this model pest.

## Materials and Methods

### Mite rearing conditions

London strain of *T. urticae* was collected from apple trees in the Vineland region in Ontario. The population was maintained for more than twenty years on beans (*Phaseolus vulgaris* ‘California Red Kidney’; Stokes, Thorold, ON) grown in soil (Pro-Mix^®^ BX Mycorrhizae™; Premier Tech, Rivière-du-Loup, QC), at 26°C under 100–150 μmol m^−2^ s^−1^ cool-white fluorescent light and a 16/8 h light/dark photoperiod.

### dsRNA synthesis

The RNeasy Mini Kit (Qiagen, Valencia, CA) and the SuperScript II cDNA Synthesis Kit (Thermo Fisher Scientific, Waltham, MA) were used for total RNA extraction and synthesis of cDNA. Template preparation for dsRNA synthesis was performed by polymerase chain reaction (PCR) amplification using specific forward and reverse primers with a minimal T7 promoter sequence at their 5’ ends (see list of primers in the Supplemental Table 1). The dsRNA-NC (382 bp) has complementarity against non-transcribed intergenic region 1690614-1690995 of the *T. urticae* genomic scaffold 12, Figure 1A. A 500 bp-chimeric dsRNA fragment was synthesized using an overlap-extension assembly PCR, resulting in the fusion between the 100-5’dsRNA-*TuVATPase* (118 bp) and the dsRNA-*NC* (382 bp) fragments. Briefly, both fragments were individually amplified using primers listed in the Supplementary Table 1. Approximately 50 ng of both PCR products (purified by Qiagen PCR purification kit) were combined and a PCR was performed for 7 cycles to combine PCR fragments into one. These PCR products were then used as a template to perform a second PCR using end primers (forward primer of *TuVATPase* and reverse primer of NC) for the final amplification of chimeric fragment. Amplified fragments were purified with the Gel/PCR DNA Fragments Extraction Kit (Geneaid Biotech, New Taipei, Taiwan) and were sequenced to confirm their identity. The RNA fragments were synthesized using 1 μg of PCR-generated fragments using the TranscriptAid T7 High Yield Transcription Kit (Thermo Fisher Scientific). Upon DNase I treatment for 30 min (Thermo Fisher Scientific), RNA was denatured (at 95°C for 5 min) and was allowed to slowly cool-down to room temperature to facilitate annealing and the formation of dsRNA. dsRNA was purified by phenol/chloroform extraction followed by ethanol precipitation. dsRNA was dissolved in nuclease-free water and quantified using a NanoDrop (Thermo Fisher Scientific, Waltham, MA).

### Preparation of ^32^P-labeled dsRNA fragments

A 1 μL aliquot of 3.3 nM ^32^P-α-UTP (Perkin Elmer) was added to the 20-μL reaction mixture of *in vitro* transcription containing 4 mM ATP, 4 mM GTP, 4 mM CTP and 0.1 mM UTP (final concentration). 2-nt 3′ overhang of dsRNAs was made using RNase T1, and DNA templates were digested by RNase-free DNase I (Takara Bio, Shiga, Japan).

### dsRNA-cleaving assay

To detect the activity of Dicer complex in mites, we used the method previously described by Fukudome *et al.*^56^and Nagano *et al.*^57^. Adult female mites (20 mg in weight) were collected with an air pump–based sucking system^58^ and were transferred to a 1.5-mL polypropylene tube. Mites were homogenized in 20 mL/g of extraction buffer containing 20 mM Tris–HCl (pH 7.5), 4 mM MgCl_2_, 5 mM DTT, 1 mM phenylmethylsulfonyl fluoride (PMSF), 1 mg/mL leupeptin and 1 mg/mL pepstatin A. Homogenates were centrifuged twice at 20,000 g for 10 to 15 min to remove debris, and the supernatant containing the protein crude extract was collected and tested for the dsRNA-cleaving activity. A 1 μL aliquot of ^32^P-labeled dsRNAs (approximately 10 ng) was incubated with 15 μL of crude protein extracts and 4 μl of 5 × dicing buffer containing 100 mM Tris–HCl (pH 7.5), 250 mM NaCl, 25 mM MgCl_2_ and 25 mM ATP with 0.1 μL of an RNase inhibitor (2313A; Takara Bio) that inhibits the enzymatic activity of RNase A but not of Dicer (RNase III), at 22°C for 2 h. Upon incubation, the cleaved products were purified by phenol/chloroform, precipitated by ethanol, separated by 15% denaturing PAGE with 8 M urea, and detected by autoradiography. The relative band intensities of the small RNA products were quantified by a Typhoon FLA 7000 image analyzer (GE Healthcare, Chicago, IL). The dsRNA-cleaving activity was calculated as the relative band intensity corresponding to the small RNAs in comparison with the total intensity of all bands in each lane. The experiment was conducted in 3 independent trials.

### dsRNA oral delivery

Fifty newly-emerged adult female mites were soaked in 50 μL of dsRNA solution (160 ng/μL, unless stated otherwise; 0.1% v/v Tween 20) at 20°C for 24 hours^34^. Subsequently, mites were washed in 100 μL of double distilled water and were transferred onto either bean leaf or bean leaf discs placed on water-soaked cotton kept at 26°C, 16/8 h light/dark photoperiod and 50% RH.

### Application of a tracer dye to monitor the uptake of dsRNA solution

Fifty newly-emerged adult female mites were soaked in 50 μL of dsRNA solution (160 ng/μL; 0.1% v/v Tween 20) mixed with 6% of blue food dye (erioglaucine; McCormick, Sparks Glencoe, MD) and were incubated at 20°C for 24 hours. After soaking, mites were washed in 100 μL of double distilled water and separated according to their color (blue or transparent) onto a bean leaf that was incubated at 26°C, 16/8 h light/dark, photoperiod and 50% RH.) The experiment was conducted in 3 independent trials. The counts of dark-body and spotless mite phenotypes upon treatment with dsRNA-*TuVATPase-*600 and dsRNA-*TuCOPB2*-B, respectively, after “blue” and “white” mite separation were analysed using the Fisher’s exact test (fisher.test function in native stats package of R). Confidence intervals of proportions were determined using Clopper-Pearson method^58^.

### Determination of mite RNAi-induced body phenotype

Two days post soaking, mite body-color associated with the application of dsRNAs was counted and frequencies calculated, except for mites treated with dsRNAs against *TuRpt3* and *TuHsc70-3* that required 4 to 5 days post dsRNA treatment for the establishment of the body-color change. A significant difference in the proportion of color phenotype, spotless, black, or normal was analyzed with Fisher’s exact test (fisher.test function in native stats package of R). For multiple comparisons of phenotype proportions between treatments Fisher’s exact test was used (fisher multcomp function, R package ‘RVAideMemoire’) and a Bonferroni correction of p-values was applied. Confidence intervals of proportions were determined using Clopper-Pearson method^58^. Images of mite body phenotypes were taken using a Canon EOS Rebel T5i camera (Canon, Japan) fitted on an upright dissecting microscope (Leica, Germany). The experiment was conducted in 3 independent trials.

### Mite survivorship

For mite survivorship, approximately 30 dsRNA treated adult female mites were placed on a detached bean leaf and mortality was recorded over 10 days. The experiment was performed in 3 independent trials. Survival curves were calculated with the Kaplan-Meier method (function survfit, R package ‘survival’) with comparisons performed based on the log-rank test (function survdiff, R package ‘survival’).

### Mite fecundity

For the treatment with dsRNA-*TuCOPB2*, the fecundity of individual adult female mites was recorded over 10 days and mites were moved every other day onto a fresh bean disc (15mm). The experiment was performed in 3 independents trials. A Wilcoxon-Mann-Whitney test was run to check the differences in fecundity between treatments (wilcox.test function in native stats package of R) with Bonferroni correction of p-values. For the fecundity assays of mites treated with dsRNAs against *Tribolium* orthologous genes, mites were recovered on a bean leaf for one-day post-treatment. The second day, 10 mites were moved from the bean leaf and placed on a bean leaf disk (15mm) and number of eggs laid by a group of 10 adult females was recorded over 3 days. Mites were moved every other day onto a fresh bean disc. The target genes were distributed between four operators and experiments were performed in 3 independent trials. All operators performed dsRNA-NC control treatment within their respective batches. Differences in the mean number of eggs laid between the gene target dsRNA and dsRNA-NC treatments per group were modelled using linear mixed-effects models with trial, treatment, and timepoint as fixed effects, using leaf-disk as the repeated measures random component and an autocorrelation structure using timepoint/leaf-disk (lme and corAR1 functions, R package ‘nlme’). The ACF function in the ‘nlme’ package was used to determine the autocorrelation for a lag of 1 in the timepoint variable. The models were validated by interrogating the qq plots of residuals (normality assumption) and ‘fitted’ versus ‘residuals’ plots to verify equal variance and linearity. The ANOVA function (R package ‘car’) was used to conduct an analysis of deviance and test hypotheses regarding fixed effects. Following a significant effect of treatment (and no confounding interactions of biological relevance), estimated marginal means and subsequent multiple comparisons between treatments were determined using the emmeans function (R package ‘emmeans’) with the application of the Tukey method of p-value adjus™ent. For plotting the mean of eggs laid per mite per day from a group of 10 mites was calculated and individual group data batches were adjusted towards an overall mean of dsRNA-NC treatment. In the box-and-whisker plots, the central line (second quartile) indicates the median, the distance between the box bottom (first quartile) and top (third quartile) indicates the interquartile range, and the whisker bottom and top indicate the minimum and maximum values, respectively.

### Real-Time Quantitative Reverse Transcription-PCR analysis of *TuCOPB2*

For the *TuCOPB2* gene expression post dsRNA treatments, 30 adult mites were collected, and RNA was extracted with the RNeasy Mini Kit and treated with DNAse (Qiagen). One microgram of RNA was reverse transcribed with the Maxima First Strand cDNA Synthesis Kit for RT-qPCR (Thermo Fisher Scientific). The reference gene *RP49* (*tetur18g03590*) coding for a ribosomal protein was used as the internal control. The RT-qPCR was performed using forward and reverse primers (shown in the Supplementary Table 1D) that amplified fragment that does not overlap by the dsRNA sequence. Cycle Threshold (Ct) values from 3 technical replicates were averaged to calculate the Ct value of each 3 independent experimental trials. Expression value for the *TuCOPB2* gene was normalized to the reference gene *RP49* and normalized quantity (NRQ) was calculated with the following equation: NRQ = (1+E_RP49_)^CtRP49^/(1+E_COPB2_)^CtCOPB2^. NRQ values were normalized to a mean of the control group treated with dsRNA-NC. Differences in the mean of normalized NRQ values between the control and treatments were analyzed with one way ANOVA followed by Dunnett’s test (function glht, R package ‘multcomp’). Differences were considered significant at *P* < 0.05 for all analyses. All analyses were performed in R (R Core Team 2015).

## Acknowledgement

This work was supported by the Government of Canada through the Ontario Research Fund (RE08-067), the Natural Sciences and Engineering Research Council of Canada (NSERC) and by the European Union’s Horizon 2020 research and innovation program (773902-SuperPests) awarded to MG and VG; and by the Japan Society for the Promotion of Science KAKENHI (Grant nos. 18H02203 to TS, 19K22304 to TF and 19K23674 to MT) and the Institute of Global Innovation Research in TUAT to MT, TS, TF and VG. MA was funded through the Global Thesis program, the University of Bari Aldo Moro, Italy.

## Author Contributions

N.B., T.S., T.F., S.G., R.N., M.G. and V.G. conceived and planned the study. N.B., S.D., M.T., M.M., D.L., M.A., P.J. and Y.A. performed experimental procedures and collected data. N.B., V.Z., T.S., K.B., M.G. and V.G. performed analysis and wrote the manuscript.

## Figure legends

**Supplemental Figure 1:**
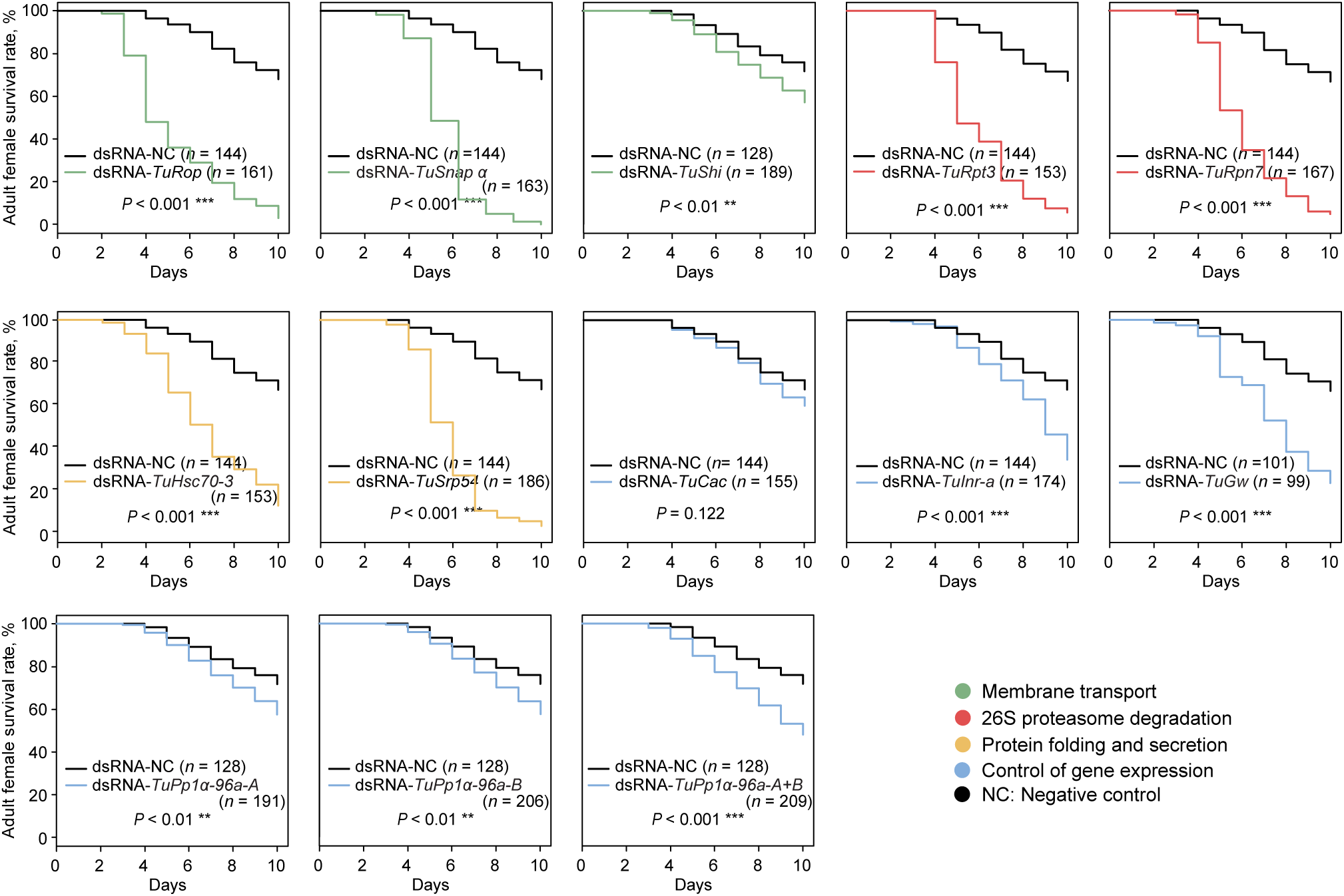
Survival curves of adult female mites after treatment with a second non-overlapping dsRNAs targeting *T. urticae* orthologs of *Tribolium* RNAi sensitive genes. Survival curves were plotted using the Kaplan-Meier method and compared using the log-rank test with Bonferroni correction (not significant, *P*>0.05; ***, *P*<0.001).

**Supplemental Table 1.**
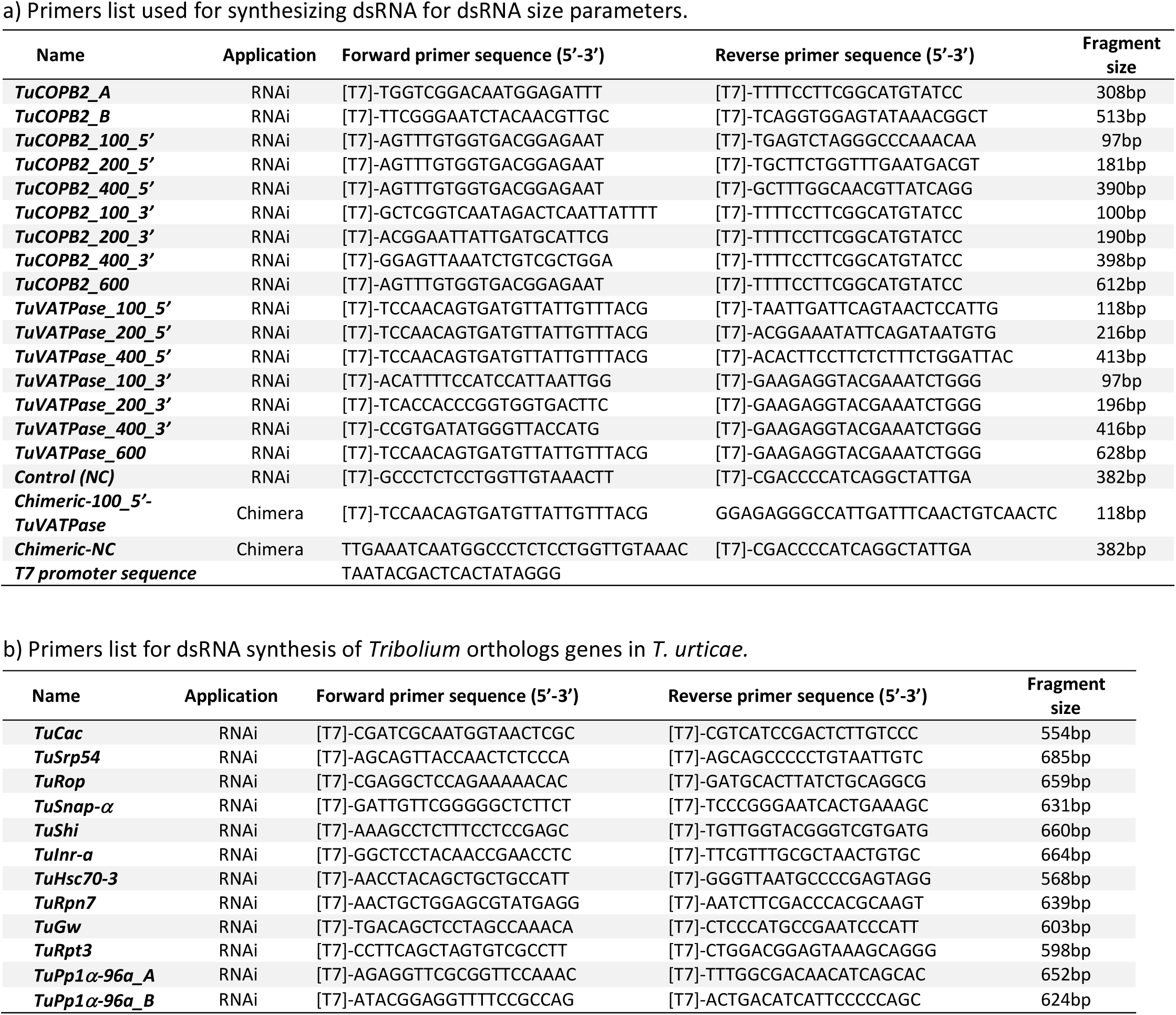
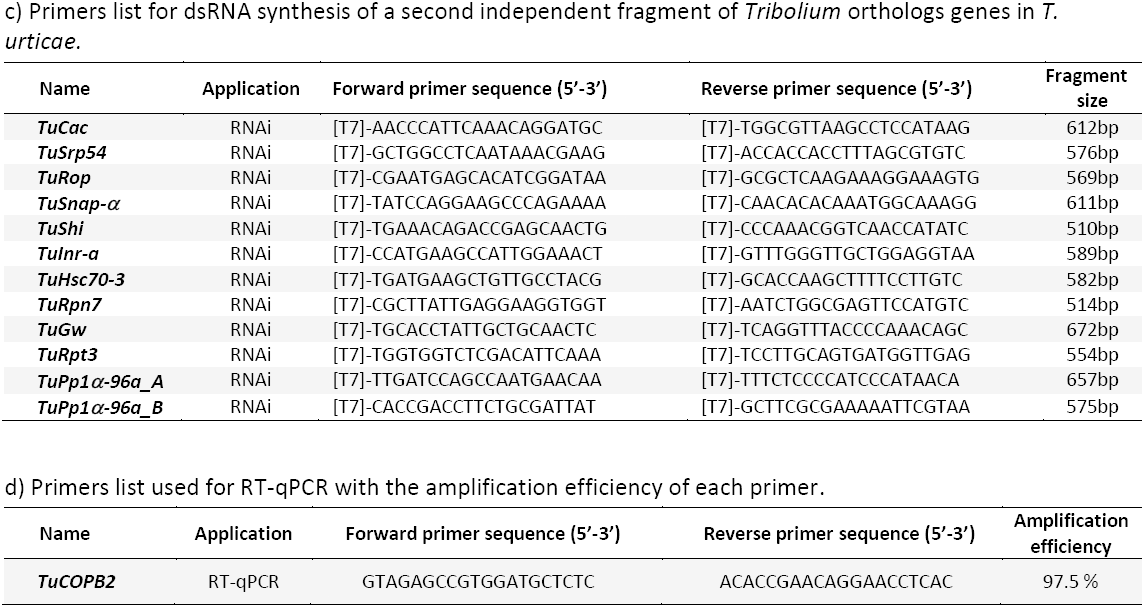
Primer list used in this study

